# Genome of *Phyllanthus emblica*: the medicinal plant Amla with super antioxidant properties

**DOI:** 10.1101/2023.05.08.539786

**Authors:** Shruti Mahajan, Manohar S. Bisht, Abhisek Chakraborty, Vineet K Sharma

**Author notes:** Correspondence: Vineet K Sharma –. **E-mail addresses of authors:** Shruti Mahajan –, Manohar S. Bisht –, Abhisek Chakraborty –, Vineet K Sharma –.

## Abstract

*Phyllanthus emblica* or Indian gooseberry, commonly known as amla, is an important medicinal horticultural plant used in traditional and modern medicines. It bears stone fruits with immense antioxidant properties due to being one of the richest natural sources of vitamin C and numerous flavonoids. This study presents the first genome sequencing of this species performed using 10x Genomics and Oxford Nanopore Technology. The draft genome assembly was 519 Mbp in size and consisted of 4,384 contigs, N50 of 597 Kbp, 98.4% BUSCO score and 37,858 coding sequences. This study also reports the genome-wide phylogeny of this species with 26 other plant species that resolved the phylogenetic position of *P. emblica*. The presence of three ascorbate biosynthesis pathways including L-galactose, galacturonate and myo-inositol pathways was confirmed in this genome. A comprehensive comparative evolutionary genomic analysis including gene family expansion/contraction and identification of multiple signatures of adaptive evolution provided evolutionary insights into ascorbate and flavonoid biosynthesis pathways and stone fruit formation through lignin biosynthesis. The availability of this genome will be beneficial for its horticultural, medicinal, dietary, and cosmetic applications and will also help in comparative genomics analysis studies.

## Introduction

Vitamin C, also known as ascorbic acid, is a vital vitamin due to its multifaceted roles in animals as well as plants, and is an essential component of the human diet (Gallie 2013; Carr and Maggini 2017). The prolonged deficiency of this vitamin causes scurvy which was infamously responsible for killing thousands of sailors in the medieval period since, humans and primates cannot synthesize vitamin C due to the absence of an enzyme gulono-lactone oxidase (GULO), which is responsible for the final conversion to ascorbic acid (Martini 2003; Wheeler, Ishikawa et al. 2015). Thus, they depend majorly on plants that are the dominant sources of vitamin C for animals (Wheeler, Ishikawa et al. 2015).

One of the richest sources of natural vitamin C is *Phyllanthus emblica*, also known as Indian gooseberry or amla. It is an economically important medicinal horticultural plant that belongs to the family Phyllanthaceae in order Malpighiales, and is widely used in pharmaceuticals, nutraceuticals, food industry, and cosmetics sectors with an estimated market of USD 49.34 billion by 2025 (Muzaffar, Sofi et al. 2022). Genus *Phyllanthus* is the largest genus of its family with approximately 1000 species of which several species are used as ethnomedicinal herbs due to the presence of medicinal phytochemicals (Sarin, Verma et al. 2014; Mao, Wu et al. 2016; Geethangili and Ding 2018). The morphological characteristics of *P. emblica* include a light grey stem with thin flaky bark, simple leaves, and greenish-yellow unisexual flowers that are arranged like female flowers on the top and male flowers on the lower side. The fruits are typical drupes of about 2 cm in diameter, also known as stone fruits, with seeds encased in a hard lignified endocarp known as stone that helps in seed protection and dispersal (Dardick and Callahan 2014; Dasaroju and Gottumukkala 2014).

The geographical distribution of this prominent ethnomedicinal herbal species spreads from tropical to warm temperate regions like India, China, Sri Lanka, Bangladesh, Indonesia, Thailand, etc., among which India is the top producers of amla with annual production of 1,275,660 metric tonnes (Mao, Wu et al. 2016; Department of Agriculture & Farmers Welfare 2021). This plant has also been used in many traditional medicine systems like Indian Ayurveda, Traditional Chinese Medicine System, etc. and is now widely used in modern medicines (Mao, Wu et al. 2016; Gul, Liu et al. 2022). Extracts of almost all parts of this plant such as leaf, bark, seed, root and fruit show medicinal properties like anti-microbial, anti-viral, anti-inflammatory, anti-oxidant, anti-aging, anti-diabetic, hypolipidemic, hypoglycaemic, neuroprotective, anti-cancer, immunomodulatory and hepatoprotective, etc. due to the presence of various secondary metabolites (phytochemicals) like alkaloids, phenolic acids, hydrolysable tannins, flavonoids, etc. with significance to human health and diseases (Gantait, Mahanta et al. 2021; Gul, Liu et al. 2022; Saini, Sharma et al. 2022; Yan, Li et al. 2022). The clinical effectiveness of *P. emblica* has been confirmed in diseases like dyslipidemia, type 2 diabetes, chronic periodontitis, symptomatic knee osteoarthritis, etc. (Gantait, Mahanta et al. 2021). Amla is used in treating COVID-19 patients where its consumption shortened the recovery time (Varnasseri, Siahpoosh et al. 2022). Additionally, its phytochemicals are reported as potential protease inhibitors of SARS-CoV-2 virus through in-silico evidences (Murugesan, Kottekad et al. 2021; Pandey, Lokhande et al. 2021). Its extracts are proven to have protective effects by maintaining gut microbiome homeostasis in vivo (Li, Lin et al. 2022; Luo, Zhang et al. 2022). Along with its benefits in human health, it is also effective in aquaculture, dairy and poultry as feed additives (Nguse, Yang et al. 2022; Van Doan, Lumsangkul et al. 2022; Abo Ghanima, Aljahdali et al. 2023).

Among the vitamin C-rich fruits, *P. emblica* is known to contain the highest content of vitamin C (up to 720mg/100g fruit) along with other phytochemicals, minerals and amino acids (Kubola, Siriamornpun et al. 2011; Chavhan 2017; Abeysuriya, Bulugahapitiya et al. 2020; Gul, Liu et al. 2022). Plants produce this vitamin to protect them against biotic (pathogens) and abiotic stresses (heat or light), and is also needed for the biosynthesis of plant hormones, and plant pigments, and acts as a cofactor in the cell cycle and metabolism, etc. (Gallie 2013). The ascorbate biosynthesis occurs in plants through four proposed pathways i.e., L-galactose (also known as Smirnoff-Wheeler pathway), galacturonate (uronic acid pathway), L-gulose and myo-inositol pathways (Fenech, Amaya et al. 2019; Paciolla, Fortunato et al. 2019). Among these pathways, the Smirnoff-Wheeler (SW) pathway is considered as the most common pathway of ascorbate biosynthesis (Gómez-García and Ochoa-Alejo 2016; Sodeyama, Nishikawa et al. 2021). Various genome-wide studies revealed ascorbic acid biosynthesis pathways in *Psidium guajava, Citrus sinensis*, kiwifruits, etc., however the ascorbate biosynthesis pathways have not been examined in *P. emblica* (Xu, Chen et al. 2013; Feng, Feng et al. 2021; Liao, Chen et al. 2021; Han, Zhang et al. 2022).

Despite being a pharmaceutically and nutritionally important plant, the genome sequence of *P. emblica* still remains unknown. However, a few transcriptome studies were carried out to explore a few biosynthesis pathways in *P. emblica* (Kumar, Kumar et al. 2016; Xiong-fang, Tai-qiang et al. 2018). Therefore, to gain genomic insights into the medicinal properties of *P. emblica*, we performed its genome sequencing and assembly using a hybrid approach that includes 10x Genomics and Oxford Nanopore Technology (ONT) long-read sequencing technologies along with transcriptomic sequencing using the Illumina technology. Further, we analysed the genes involved in vitamin C, lignin and flavonoid biosynthesis pathways. We also constructed a genome-wide phylogenetic tree of *P. emblica* with 26 plant species, which were further analysed for gene family expansion and contraction. Furthermore, this study performed a comprehensive comparative evolutionary genomic analysis across 19 plant species to uncover the genes with multiple signatures of adaptive evolution in *P. emblica*.

## Materials and Methods

### DNA-RNA extraction, species identification and sequencing

The leaves sample from a plant was collected from Bhopal, India (23.2599° N, 77.4126° E). The DNA sample extracted from the leaves sample using Carlson lysis buffer except for the precipitation step that was carried out with 0.5X volume of NaCl ad 0.7X volume of isopropanol (Jaiswal, Mahajan et al. 2021). The extracted DNA was quantified, and quality was checked on Qubit 2.0 fluorometer and Nanodrop 8000 spectrophotometer, respectively. Species identification assay was performed using marker genes: Internal Transcribed Spacer (ITS) and Maturase K (MatK). The extracted DNA was utilized to prepare libraries for 10x Genomics and nanopore sequencing that were sequenced on Illumina NovaSeq 6000 and MinION Mk1C sequencers, respectively. The RNA extraction was performed as per the protocol described by Kumar and Singh (2012) with a few modifications (Kumar and Singh 2012). The RNA was used for preparing the library using TruSeq Stranded Total RNA Library Preparation kit (Illumina Inc., CA, USA) with Ribo-zero Plant workflow and sequenced on Illumina NovaSeq 6000 instrument for generated 150 bp paired-end reads.

### Genome assembly

The proc10xG set of python scripts (https://github.com/ucdavis-bioinformatics/proc10xG) was used to pre-process the 10x Genomics raw reads by removing the barcode sequences. The obtained reads were processed by SGA-preqc (paired-end mode) for genome size estimation which works on a k-mer distribution-based approach (Simpson and Durbin 2012). For genome complexity assessment, these pre-processed reads were used by Jellyfish v2.2.10 and GenomeScope v2.0 for generating k-mer count histograms and calculating heterozygosity, respectively (Marçais and Kingsford 2011; Ranallo-Benavidez, Jaron et al. 2020).

Guppy v3.2.1 (Oxford Nanopore Technologies) was used to carry out base calling of Nanopore raw reads. Adaptor removal was performed on this base called raw data using Porechop v0.2.4 (Oxford Nanopore Technologies). The pre-processed reads were utilized for genome assembly using three different assemblers: wtdbg v2.0.0, SMARTdenovo (https://github.com/ruanjue/smartdenovo), and Flye v2.9 (Kolmogorov, Yuan et al. 2019; Ruan and Li 2020). wtdbg v2.0.0 and Flye v2.9 were used with default settings whereas SMARTdenovo was used with 0 as minimum read length (Kolmogorov, Yuan et al. 2019; Ruan and Li 2020). Quast v5.0.2 was employed to assess the genome assembly statistics (Gurevich, Saveliev et al. 2013). The genome assembly resulting from Flye v2.9 was considered for further analysis due to its better assembly statistics and assembled genome size (Kolmogorov, Yuan et al. 2019). The assembly was polished three times by Pilon v1.24 using filtered reads. ARCS v1.2.2 and LINKS v2.0.0 (default settings) were employed for the first round of scaffolding using Longranger basic v2.2.2 (https://support.10xgenomics.com/genome-exome/software/pipelines/latest/installation) barcode filtered 10x Genomics linked reads and adaptor-removed Nanopore reads, respectively (Walker, Abeel et al. 2014; Warren, Yang et al. 2015; Yeo, Coombe et al. 2018). After scaffolding, the quality-filtering of RNA-Seq paired-end raw reads was performed using Trimmomatic v0.39 with parameters-“LEADING:15 TRAILING:15 SLIDINGWINDOW:4:15 MINLEN:50” which were subsequently utilized by AGOUTI v0.3.3 for the second round of scaffolding (Bolger, Lohse et al. 2014; Zhang, Zhuo et al. 2016).

Supernova v2.1.1 was used to perform *de novo* assembly of *P. emblica* with maxreads =all options with other default parameters (Weisenfeld, Kumar et al. 2017). The obtained genome assembly was corrected by Tigmint v1.2.6 using Longranger basic v2.2.2 processed linked reads (Jackman, Coombe et al. 2018). Further, the first round of scaffolding was performed with ARCS v1.2.2 and LINKS v2.0.0 using Longranger basic v2.2.2 processed linked reads and adaptor removed Nanopore reads, respectively (Warren, Yang et al. 2015; Yeo, Coombe et al. 2018). To increase the assembly contiguity, AGOUTI v0.3.3 was performed using the pre-processed transcriptomic paired-end reads (quality-filtered) (Zhang, Zhuo et al. 2016).

RagTag v2.1.0 was used to merge both the assemblies obtained from Supernova and Flye assemblers using the patch command line utility (Alonge, Lebeigle et al. 2021). RagTag will use reference as the main assembly and use query assembly to fill the gap in the reference assembly. LR_Gapcloser was used to perform gap-closing of the assembly using pre-processed Nanopore reads (Xu, Xu et al. 2019). Sealer v2.3.5 was used for gap-closing of the assembly using barcode-removed linked reads with 30-120 k-mer value and 10 bp interval (Paulino, Warren et al. 2015). The fixation of small indels, base errors, and local misassemblies was performed by Pilon v1.24 using pre-processed linked reads to provide the draft genome assembly of *P. emblica* (Walker, Abeel et al. 2014). Obtained draft genome assembly was further length base filtered and scaffolds having length ≥5 kbp were retained. BUSCO v5.2.2 with embryophyta_odb10 single-copy orthologs dataset was employed to assess the completeness of genome assembly (Simão, Waterhouse et al. 2015). For further assessment of the assembly quality, the barcode-filtered 10x Genomics reads, nanopore long reads and the quality-filtered transcriptomic reads were mapped onto the genome assembly using BWA-MEM v0.7.17, MiniMap2 v2.17 and HISAT2 v2.2.1, respectively, and SAMtools v1.13 “flagstat” was used to calculate the percentage of mapped reads (Li, Handsaker et al. 2009; Li 2013; Kim, Langmead et al. 2015; Li 2018).

### Annotation of genome and construction of gene set

For annotation of repeats in final genome assembly, RepeatModeler v2.0.3 was used to generate a *de novo* repeat library (Sharma, B-Rao et al. 2003; Sharma, Brahmachari et al. 2005; Flynn, Hubley et al. 2020). The clustering of repeats was performed using CD-HIT-EST v4.8.1 with parameters - 8 bp seed size and 90% sequence identity to eliminate redundant sequences (Fu, Niu et al. 2012). The resultant repeat library was utilized by RepeatMasker v4.1.2 (http://www.repeatmasker.org) to soft-mask the final genome assembly of *P. emblica*.

The coding gene set was constructed on the resultant repeat-masked genome assembly using MAKER v3.01.04 pipeline which deploys approaches such as *ab initio* and evidence alignment for prediction (Campbell, Holt et al. 2014). The construction of *de novo* transcriptome assembly was performed by Trinity v2.14.0 (default parameters) using quality-filtered transcriptomic reads of *P. emblica* from this study and previously reported studies (Haas, Papanicolaou et al. 2013; Liu, Ma et al. 2018). The gene set was constructed with transcriptome assembly and protein sequences of species belonging to the same order Malpighiales (*Populus trichocarpa* and *Manihot esculenta*) that were used as EST and protein evidence, respectively. In the MAKER pipeline, AUGUSTUS v3.2.3 was used for *ab initio* gene prediction while empirical evidence alignments and alignment polishing were performed using BLAST and Exonerate v2.2.0 (https://github.com/nathanweeks/exonerate), respectively (Altschul, Gish et al. 1990; Stanke, Keller et al. 2006). Based on the length and Annotation Edit Distance (AED) of gene models, the final gene set was constructed by selecting genes with length (≥150bp) and AED values <0.5. The completeness of this final gene set (also termed a high-confidence gene set) was checked using BUSCO v5.2.2 through the embryophyta_odb10 dataset (Simão, Waterhouse et al. 2015).

Additionally, Barrnap v0.9 (https://github.com/tseemann/barrnap) and tRNAscan-SE v2.0.9 were used to perform *de novo* prediction of rRNA and tRNA, respectively (Chan and Lowe 2019). Based on homology, miRNA gene sequences in the *P. emblica* genome were identified using miRbase database with e-value 10^−9^ and 80% identity (Griffiths-Jones, Saini et al. 2007).

### Phylogenetic tree construction

The 26 plant species were selected from Ensembl plant release 54 for phylogenetic analysis considering the representation of each plant family among the selected species (Bolser, Staines et al. 2016). Besides the protein sequences of these selected 25 eudicots and one monocot species, MAKER-derived protein sequences of *P. emblica* were used for phylogenetic tree construction. Among all the protein files, the longest isoform for each protein was selected and provided to OrthoFinder v2.5.4 to construct the set of orthologous genes (Emms and Kelly 2019). KinFin v1.0 was used to extract fuzzy one-to-one orthologs protein sequences that were present in all 27 species (Laetsch and Blaxter 2017). MAFFT v7.310 was used to individually align all the obtained fuzzy one-to-one orthologs which were filtered and concatenated using BeforePhylo v0.9.0 (https://github.com/qiyunzhu/BeforePhylo) (Katoh and Standley 2013). These obtained protein sequences were used to construct a phylogenetic tree based on maximum likelihood using RAxML with the ‘PROTGAMMAAUTO’ amino acid substitution model and 100 bootstrap values (Stamatakis 2014).

### Gene family expansion and contraction analysis

The proteome files containing the longest isoform for every protein from selected 27 species along with generated species phylogenetic tree were provided to CAFE v5 to assess the evolution of gene families (De Bie, Cristianini et al. 2006). The species phylogenetic tree was adjusted to an ultrametric tree based on the calibration point of 120 million years between *P. emblica* and *Beta vulgaris* obtained from TimeTree database v5.0 (Kumar, Suleski et al. 2022). BLASTP was performed on protein sequences of all 27 species in All-versus-All mode (Altschul, Gish et al. 1990). The BLASTP results were clustered using MCL v14-137 and gene families containing clade-specific genes and >100 gene copies for minimum of one species were eliminated. These resultant gene families and ultrametric species tree were used to analyse the evolution (expansion/contraction) of gene families using a two-lambda (λ) model where λ signifies a random birth-death parameter. Among the obtained contracted and expanded gene families, gene families with >10 genes were considered as highly expanded gene families.

### Identification of signatures of adaptive evolution

The 19 plant species were selected for identification of genes with evolutionary signatures that included five species of order Malpighiales i.e., *Linum usitatissimum, Manihot esculenta, Phyllanthus emblica, Populus trichocarpa* and *Ricinus communis* along with *Actinidia chinensis* (order Ericales), *Arabidopsis thaliana* (order Brassicales), *Coffea canephora* (order Gentianales), *Cucumis sativus* (order Cucurbitales), *Daucus carota* (order Apiales), *Eucalyptus grandis* (order Myrtales), *Ficus carica* (order Rosales), *Gossypium raimondii* (order Malvales), *Helianthus annuus* (order Asterales), *Olea europaea* (order Lamiales), *Pistacia vera* (order Sapindales), *Quercus lobata* (order Fagales), *Solanum lycopersicum* (order Solanales) and *Vitis vinifera* (order Vitales). Orthologous gene sets were constructed by Orthofinder v2.5.4 using the proteome files from 19 selected plant species. Orthogroups that contained protein sequences from all these selected species were retrieved and in case multiple protein sequences were present for a species, the longest isoform of that protein was selected and retained for further analysis.

### Identification of genes with higher rate of evolution

The resulting orthogroups across 19 plant species were aligned individually using MAFFT v7.310 (Katoh and Standley 2013). These obtained alignments were used to construct a phylogenetic tree for individual orthogroups using RAxML v8.2.12 with ‘PROTGAMMAAUTO’ amino acid substitution model and a bootstrap value of 100 (Stamatakis 2014). R package “adephylo” was used to calculate root-to-tip branch length distance for genes of all species in the phylogenetic trees. The genes of *P. emblica* with comparatively higher root-to-tip branch length distance values were extracted and listed as the genes with higher nucleotide divergence or rate of evolution.

### Identification of *P. emblica* genes with unique amino acid substitutions

Using the multiple sequence alignments obtained from MAFFT v7.310, which were used for the identification of genes with a high rate of evolution, amino acid positions alike in all the species except *P. emblica* were extracted and labelled as genes with unique amino acid substitution. Ten amino acids around any gap were not included in this analysis. The functional impact of obtained genes showing amino acid substitution was evaluated using Sorting Intolerant From Tolerant (SIFT) with UniProt database (Ng and Henikoff 2003).

### Identification of positively selected genes

MAFFT v7.310 was used for individual alignment of nucleotide sequence of all orthologous gene sets across selected 19 species (Katoh and Standley 2013). PAML v4.9a with “codeml” program based on a branch-site model used nucleotide alignments in PHYLIP format and a species phylogenetic tree of these 19 species (constructed using RAxML) to identify genes with positive selection (Yang 2007). These obtained genes with their log likelihood values were further processed through likelihood-ratio tests and genes with FDR-corrected p values of <0.05 were labelled as positively selected genes. Positively selected codon sites were identified using Bayes Empirical Bayes (BEB) analysis with criteria of >95% probability for the foreground lineage.

### Genes with multiple signatures of adaptive evolution (MSA)

The high rate of evolution, unique amino acid substitution with functional impact and positive selection are the evolutionary signatures of adaptive evolution. *P. emblica* genes that showed at least two of these evolutionary signatures were considered as the genes with multiple signatures of adaptive evolution (MSA) (Jaiswal, Gupta et al. 2018; Mittal, Jaiswal et al. 2019; Chakraborty, Mahajan et al. 2021; Jaiswal, Mahajan et al. 2021; Chakraborty, Mahajan et al. 2022).

### Functional annotation

The annotation of high-confidence gene sets of *P. emblica* was performed using NCBI non-redundant (nr) database, SWISS-PROT database and Pfam-A v32.0 database using BLASTP (10^−5^ e-value), BLASTP (10^−5^ e-value), and HMMER v3.3, respectively (Bairoch and Apweiler 2000; Bateman, Coin et al. 2004; Finn, Clements et al. 2011; Sharma, Gupta et al. 2015). The considered contracted and expanded gene families of *P. emblica* were extracted and provided to KAAS v2.1 and eggNOG mapper v2.1.9 for functional annotation, respectively (Moriya, Itoh et al. 2007; Huerta-Cepas, Forslund et al. 2017). The functional annotation of contracted and expanded gene families was also checked manually on Kyoto Encyclopedia of Genes and Genomes (KEGG) database. The KEGG Orthology (KO) and KEGG pathway assignment of gene set from comparative evolutionary analysis were also carried out using KAAS v2.1 genome annotation server and eggNOG-mapper v2.19 (Moriya, Itoh et al. 2007; Huerta-Cepas, Forslund et al. 2017).

### Analysis of vitamin C biosynthesis genes

The protein sequences of all the available enzymes of all four proposed pathways of vitamin C biosynthesis for *A. thaliana* were downloaded from Swiss-Prot or NCBI database. The gene D-galactose reductase (*GalUR*) was not available for *A. thaliana*, thus, sequence from strawberry plant species was used. These protein sequences were matched against the protein sequences of *P. emblica* using BLASTP with e-value 10^−9^ (Altschul, Gish et al. 1990). The enzymes involved in galactose pathway from vitamin C-rich plants i.e., *Actinidia chinensis* (kiwi), *Capsicum annuum* (chilli pepper), *Carica papaya* (papaya), *Citrus sinensis* (sweet orange), *Malpighia glabra* (acerola), *Myrciaria dubia* (camu-camu), *Solanum lycopersicum* (tomato) and *Vitis vinifera* (grapes) along with Arabidopsis thaliana as an outgroup were also obtained from UniProt or NCBI databases. Six genes i.e., Mannose 1-phosphate guanylyl transferase (*GMPP*), GDP-D-Mannose 3’,5’-epimerase (*GME*), GDP-L-galactose phosphorylase (*GGP/VTC2/VTC5*) and L-galactose-1-phosphate phosphatase (*GPP/VTC4*), L-galactose dehydrogenase (*GalDH*) and L-galactono-1,4-lactone dehydrogenase (*GLDH*) were selected for phylogeny due to their sequence availability for all nine species. The phylogenetic tree was constructed using these six genes each from 10 selected species (including *P. emblica*) using RAxML (Stamatakis 2014). Among the above expanded and contracted genes, we checked for the copy number of all the gene families involved in ascorbate biosynthesis and regeneration pathways in the *P. emblica* genome.

### Analysis of Flavonoid biosynthesis pathway

The protein sequences of genes involved in flavonoid biosynthesis pathway for *Manihot esculenta* were downloaded from UniProt and NCBI databases and matched against the protein sequences of *P. emblica* using BLASTP (e-value 10^−9^) (Altschul, Gish et al. 1990).

### Analysis of Lignin biosynthesis pathway

The protein sequences of genes involved in lignin biosynthesis pathway for *Arabidopsis thaliana* were downloaded from UniProt and NCBI databases and matched against the protein sequences of *P. emblica* using BLASTP (e-value 10^−9^) (Altschul, Gish et al. 1990).

## Results

### Species identification and sequencing

The species identification was performed using ITS and MatK DNA markers that aligned to *P. emblica* sequences with the highest identity of 100% and 99.89%, respectively, and confirmed the species. A total of 136 Gbp (∼237x coverage) and 18.3 Gbp (∼32x coverage) of genome sequence data were generated using third-generation sequencing technologies i.e., 10x Genomics and Oxford Nanopore Technology (ONT), respectively. Further, ∼85 million transcriptomic reads from leaf tissue were used for analysis in this study.

### Genome assembly and annotation

We computationally estimated the genome size of *P. emblica* to be 572 Mbp. The final genome assembly had a size of 519 Mbp and consisted of 4,384 contigs with GC content of 33.49%, largest contig of 3.3 Mbp, and N50 of 597 Kbp. The heterozygosity was estimated to be 1.37%, which appears to be high given its small genome size. The 98.4% complete and 0.4% fragmented BUSCOs of this genome assembly indicated its completeness. Further, 96.8% of linked reads and 93.45% of nanopore long reads could be mapped on the final genome assembly. The repeats constituted 53.39% of the genome with 2,051 *de novo* repeat family sequences that clustered into 1,803 repeat families. Among the interspersed repeats, 10.96% and 10.13% were predicted as Ty1/Copia and Gypsy/DIRS1 elements, respectively. A total of 216 microRNA (miRNA), 815 transfer RNA (tRNA) and 141 ribosomal RNA (rRNA) genes were identified in the genome.

The *de novo* transcriptome assembly comprised of a total of 238,454 transcripts and these transcripts were used as EST (empirical evidence) in the MAKER pipeline. The high-confidence gene set constituted of 37,858 genes that were obtained by filtering the gene models based on length ≥150 and AED value<0.5. This high-confidence gene set had an 89.9% complete BUSCO score. Overall, ∼ 96% (36,296 out of 37,858) high-confidence coding genes of *P. emblica* could be annotated using the three reference databases; NCBI-nr, Swiss-Prot, and Pfam-A.

### Phylogenetic tree construction

OrthoFinder utilized the protein sequences from 27 plant species and identified 145,194 orthogroups, of which KinFin predicted 123 one-to-one fuzzy orthogroups. The 123 one-to-one fuzzy orthogroups were aligned, and these alignments were concatenated, and selected based on the absence of any undetermined values. These selected concatenated protein sequence alignments containing 104,108 alignment positions were used to construct a phylogenetic tree based on maximum likelihood using 26 eudicot species and *Zea mays* as an outgroup. The phylogenetic tree indicated *Populus trichocarpa* and *Manihot esculenta* as the closest species to *P. emblica* as they belong to the same order Malpighiales. As per the phylogenetic tree, *P. emblica* diverged earlier than the other considered species of the order Malpighiales (**Fig. 1**). Similarly, the phylogeny constructed with vitamin C biosynthesis genes followed the genome-wide phylogeny of *P. emblica* where the species like *P. emblica* and *Malpighia glabra, Arabidopsis thaliana* and *Carica papaya* and, *Capsicum annuum* and *Solanum lycopersicum* belonging to the same orders were sharing their common ancestral node (**Fig. 2**).

**Figure 1.**
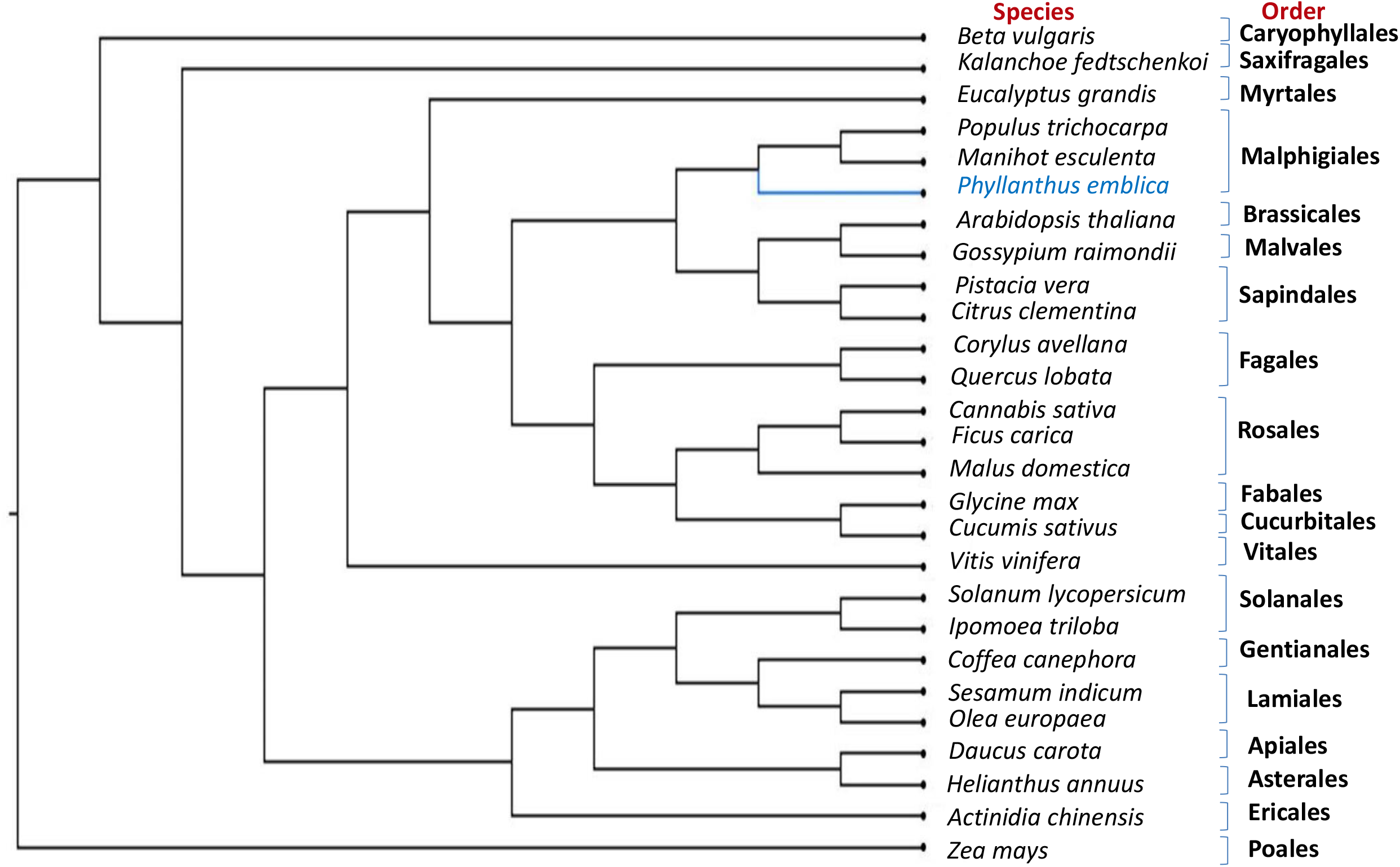
Genome-wide phylogeny of *P. emblica* with 26 other plant species. Genome-wide phylogeny of *P. emblica* with 25 other eudicot species and a monocot species, *Zea mays* that was used as an outgroup.

**Figure 2.**
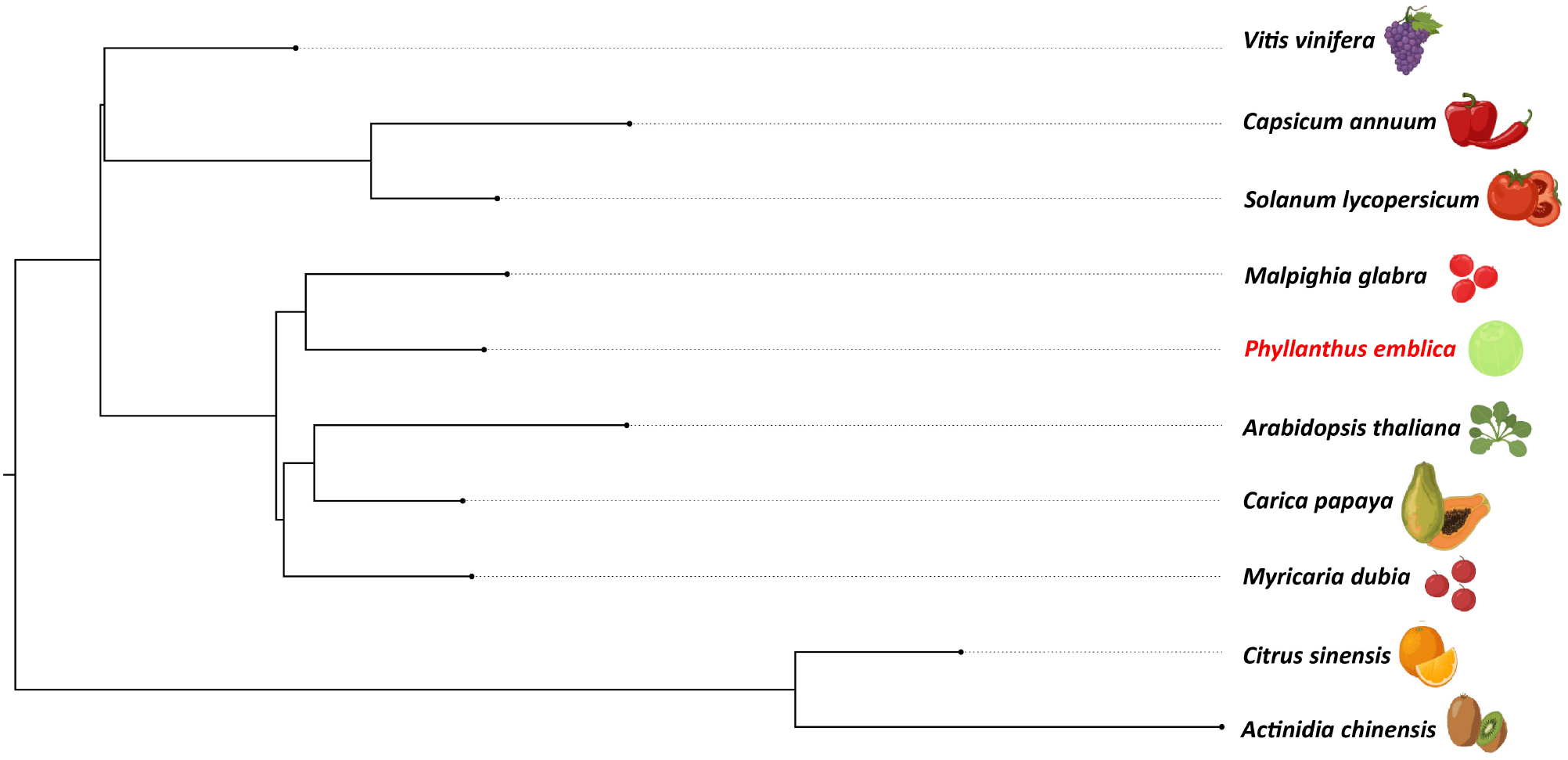
Vitamin C biosynthesis genes phylogeny of *P. emblica* with other vitamin C-rich fruits. The phylogeny constructed for *P. emblica, Actinidia chinensis, Capsicum annuum, Carica papaya, Citrus sinensis, Myrciaria dubia, Malpighia glabra, Solanum lycopersicum, Vitis vinifera* and *Arabidopsis thaliana* using six genes of Smirnoff Wheeler pathway of ascorbate biosynthesis. The genes used were GDP-mannose pyrophosphorylase (*GMPP*), GDP-D-Mannose 3’,5’-epimerase (*GME*), GDP-L-galactose phosphorylase (*GPP/VTC2*), L-galactose-1-phosphate phosphatase (GPP/*VTC4*), L-galactose dehydrogenase (*GalDH*) and L-galactono-1,4-lactone dehydrogenase (*GLDH*).

### Gene family expansion and contraction analysis

Gene family expansion and contraction analysis helps to identify the gene families that have increased or decreased in number in a given species. The adaptive evolution analysis provides the clues of natural selection of specific phenotypic traits in a given species to cope with diverse environmental conditions that help in its survival. CAFÉ analysis showed a contraction of 1,048, and an expansion of 3,520 gene families in this species. Among these families, five and 42 gene families were found to be highly contracted and highly expanded, respectively. Among the 42 expanded gene families, 38 could be functionally annotated. The expanded gene families were majorly involved in lignin biosynthesis pathway, MAPK signalling pathway, transcription (as transcription factors), phenylpropanoid pathway, brassinosteroid biosynthesis, terpenoid biosynthesis, transportation (as transporters), plant hormone signal transduction, etc.

### Identification of signatures of adaptive evolution

For evolutionary analysis, 7,864 orthogroups were obtained from MAFFT across 19 selected species. Among these orthogroups, 46 genes showed higher nucleotide divergence and 488 genes showed positive selection in *P. emblica*. Approximately 35% (2,791 of 7,864) of orthogroups showed unique amino acid substitutions. The genes that show any two of the three signatures of adaptive evolution i.e., high rate of nucleotide divergence, positive selection and unique amino acid substitutions were considered as genes with multiple signatures of adaptive evolution (MSA). A total of 236 genes were identified as MSA genes for *P. emblica*. The MSA genes were found to be involved in physiological processes like plant growth, ROS regulation and detoxification, DNA damage response, immune signalling, abiotic stress response, pathogen resistance and response, response to hormones like ethylene, abscisic acid, gibberellin and cytokinin, and cell wall modification.

### Vitamin C biosynthesis pathway

Ascorbate, a non-enzymatic antioxidant, plays an important role in ROS detoxification and is a part of the ascorbate-glutathione pathway. *P. emblica* contains all the genes of the SW pathway similar to the other vitamin C-rich plant species like guava, kiwi, chilli pepper, etc., (Gómez-García and Ochoa-Alejo 2016; Wang, Yu et al. 2018; Feng, Feng et al. 2021). The phylogeny of six genes of SW pathway showed closeness of *P. emblica* with *Malpighia glabra*, which also lies in the same order Malphigiales (**Fig. 2**). Further, the genome clues for presence of another pathway of ascorbate biosynthesis, i.e. galacturonate pathway in *P. emblica* genome. This pathway is also proposed in tomatoes, strawberries, oranges, and grapes due to the presence of the key gene of this pathway and was also present in *P. emblica* (Agius, González-Lamothe et al. 2003; Cruz-Rus, Botella et al. 2010; Cruz-Rus, Amaya et al. 2011; Badejo, Wada et al. 2012; Xu, Chen et al. 2013). In addition, the genes involved in myo-inositol pathway were found, which supports the presence of the third pathway of ascorbate biosynthesis in *P. emblica*. However, the presence of the fourth pathway (L-gulose pathway) could not be confirmed due to lack of sufficient identification of genes involved in this pathway in this genome.

Among the genes of all proposed ascorbate biosynthesis pathways, six genes were identified with unique amino acid substitutions. Genes associated with galacturonate pathway of ascorbate biosynthesis were found with MSA, unique amino acid substitution and highly expanded.

### Glutathione metabolism and Ascorbate-glutathione pathway

Glutathione is another non-enzymatic antioxidant that plays a key role in different environmental stresses mainly oxidative stress and is a part of the ascorbate-glutathione pathway (Hasanuzzaman, Bhuyan et al. 2019; Dorion, Ouellet et al. 2021). Genes involved in glutathione metabolism showed positive selection and multiple signatures of adaptive evolution.

The ascorbate-glutathione pathway, also known as ascorbate regeneration cycle, plays an important role in oxidative stress response by converting the oxidised ascorbate forms to ascorbate via four enzymes i.e., ascorbate oxidase (AO), ascorbate peroxidase (APX), dehydroascorbate reductase (DHAR) and monodehydroascorbate reductase (MDHAR) (Chen, Young et al. 2003; Li, Huang et al. 2017). All these genes of ascorbate regeneration pathway were found in *P. emblica*. Among the four enzymes, three genes showed multiple signatures of adaptive evolution and unique amino acid substitutions.

### Flavonoid biosynthesis pathway

All 15 key genes of flavonoid biosynthesis pathway were found in *P. emblica* genome, and seven genes of these genes contained unique amino acid substitutions. Six genes were highly expanded and among which one gene showed multiple signatures of adaptation.

### Lignified endocarp in this stone fruit

A lignified endocarp is a trait found in stone fruits like *P. emblica*. Lignin is important in stone cell formation in a drupe fruit that provides rigidity for seed protection and dispersal. Eight out of 13 genes of lignin biosynthesis were identified with unique amino acid substitutions. Among these eight genes, one enzyme gene showed multiple signatures of adaptive evolution. Furthermore, six gene families were highly expanded. Moreover, the gene families of two transcription factors are involved in lignin biosynthesis pathway were also found to be highly expanded.

### Plant growth, hormone and stress response

Among the 236 MSA genes, 36 genes were found to be involved in plant growth and development. These 36 genes are involved in cell division, flower development, seed development, seed germination, shoot development, cell elongation, sugar metabolism, cell wall biosynthesis, and root development, etc. Phytohormones such as auxin, cytokinin, gibberellic acid, abscisic acid (ABA), ethylene, etc. help in plant growth, development, stress tolerance, etc. throughout the plant life. Twenty MSA genes were responsible for plant hormone biosynthesis, signalling and response in plants. These include genes associated with plant hormone responses related to auxin, abscisic acid, ethylene, jasmonic acid, and cytokinin. Plants have mechanisms for stress response against abiotic and biotic stresses. 38 out of the 236 MSA genes were associated with various responses against these stresses. Among these, 30 genes were associated with responses to abiotic stresses like salt, cold, heat, drought, etc., whereas 19 genes were associated with biotic stress tolerance. These genes are involved in stress responses like stress signal transduction, secondary metabolite biosynthesis, abscisic acid biosynthesis, ROS detoxification, stress-specific gene regulation, degradation of misfolded and damaged proteins, DNA damage response, cell wall modification, disease resistance, etc. Also, six MSA genes were found to be involved in DNA damage repair mechanism in plants against plant stresses.

### ROS regulation and detoxification

Reactive oxygen species (ROS) are metabolic by-products produced in mitochondria, plastids and peroxisomes, which can cause irreversible DNA damage resulting into cell death. In plants, ROS not only cause harmful effects but also act as signalling molecules for plant growth and stress responses. Eighteen out of the 236 MSA genes were associated with ROS response, regulation and detoxification. These genes are involved in porphyrin biosynthesis, ROS-induced responses, ROS scavenging, biosynthesis and protecting antioxidant enzymes, maintaining homeostasis, accumulation, and biosynthesis of antioxidants, and activation of Fe-S cluster.

## Discussion

*P. emblica* or amla is a widely used medicinal plant with enormous antioxidant properties (Gul, Liu et al. 2022). To understand the genomic basis of these properties, we successfully built the first draft genome assembly of *P. emblica* using a hybrid sequencing approach involving Illumina-10x Genomics and ONT technologies. The implementation of hybrid approach helped in constructing a high-quality genome assembly of this repetitive and highly heterozygous genome. Further, this study is the first to resolve the genome-wide phylogenetic position of *P. emblica* with respect to 26 other plant species and found its early divergence from *Manihot esculenta* and *Populus trichocarpa* species of order Malpighiales. Our phylogeny is also supported by the revised classification of order Malpighiales where family Phyllanthaceae was separated as individual family from family Euphorbiaceae in the Angiosperm Phylogeny Group Classification (APG III) (Group 2009; Kawakita and Kato 2017). The phylogeny of vitamin C biosynthesis genes also followed the genome-wide phylogeny. Further, key genomic insights were gained from the results of gene family expansion and contraction and from the genes with multiple signatures of adaptive evolution in *P. emblica*. The genes related to the biosynthesis of ascorbic acid, lignin and flavonoid were found to be evolutionarily selected in *P. emblica*.

Ascorbic acid is the major antioxidant in *P. emblica* and its fruit “amla” is one of it richest natural sources. A transcriptomic study of oranges had shown the attribution of different pathways of ascorbate biosynthesis in a tissue-specific as well as fruit developmental stage specific manner (Caruso, Russo et al. 2021). This could also be possible in *P. emblica* where the presence of three pathways of ascorbate biosynthesis is traced, and they could have roles in different stages and tissues. It was noted that the genes of one of the ascorbic acid biosynthesis pathways i.e., galacturonate pathway were found with MSA, amino acid substitutions and highly expanded gene family in the *P. emblica* genome. The involvement of these enzymes in increased ascorbate production in tomatoes along with their role in regulating ascorbate content through galacturonate pathway is shown in previous studies (Di Matteo, Sacco et al. 2010; Badejo, Wada et al. 2012; Ruggieri, Sacco et al. 2015; Rigano, Lionetti et al. 2018). Thus, it is tempting to speculate that the evolution of genes of galacturonate pathway could be associated with the high ascorbate production in *P. emblica*.

*P. emblica* is also rich in flavonoids that are synthesized in response to plant abiotic stress and contribute to its antioxidant property. The genes involved in the initial part of flavonoid biosynthesis pathway were found with unique amino acid substitutions, and the gene that showed multiple signatures of adaptation is previously reported to increase flavonoid accumulation (Wang, Xu et al. 2014; Nguyen, Hoang et al. 2021). In addition to these, six gene families were highly expanded which collectively indicates evolution of flavonoid biosynthesis genes in *P. emblica*. These genes are involved in biosynthesis of flavonoids such as isoflavones, flavones, anthocyanins and flavonols that have antioxidant properties and provide tolerance to various abiotic and biotic stresses (Verhoeyen, Bovy et al. 2002; Agati, Azzarello et al. 2012). The evolutionary selection of these flavonoid associated genes might be responsible for the high antioxidant property and stress tolerance of *P. emblica*.

Being a stone fruit, a lignified endocarp is a trait found in the fruits of *P. emblica*, and the evolution of lignin biosynthesis pathway was thus one of the major findings. Lignin is important for the stony seed coat formation in drupes that provides rigidity for its protection. The lignin biosynthesis genes were observed to be highly expanded and among the MSA genes in *P. emblica* genome, which hints towards the evolutionary significance of lignified endocarp in this stone fruit. The lignin biosynthesis gene families were also reported to be expanded in the other stone fruit genomes such as pear and *Populus*, which are economically important due to their fruit and wood, respectively, where lignin is the main content of pear’s stone cells and poplar’s wood (Wu, Wang et al. 2013; Li, Xing et al. 2022).

*P. emblica* also produces a large variety of secondary metabolites that provide tolerance against plant abiotic stresses. The expansion of gene families and MSA in genes involved in biosynthesis of various secondary metabolites and pathogen resistance against abiotic and biotic stresses, respectively were found that indicates the evolution of stress tolerance genes in this genome. In addition to plant stress tolerance, the genes involved in ROS regulation and detoxification were also evolved in *P. emblica*.

Taken together, it is apparent that the adaptive evolution in genes involved in ascorbate biosynthesis, glutathione metabolism, flavonoid biosynthesis, and ROS detoxification are associated with the high antioxidant potential of *P. emblica*, which makes it a valuable herbal plant for use in traditional and modern medicine, horticulture, food and cosmetic products. Further, the high concentration of vitamin C in the amla fruit and the large produce (upto 100 kg) of fruits per tree compared to other vitamin C rich fruits like *Malpighia glabra* (15-30 Kg/tree) and *Myrciaria dubia* (25-30 Kg/tree), makes it the perfect choice in switching from synthetic to natural supplementation of vitamin C (Rodrigues, De Menezes et al. 2001; Orwa, Mutua et al. 2009; Carr and Vissers 2013). Further, this plant also shows high genetic diversity and easy adaptation to various climatic zones and environmental conditions (Liu, Ma et al. 2020). The availability of the first draft genome of this economically important plant is likely to help in developing improved nutraceuticals, food, cosmetics and pharmaceutical products, and for further horticultural and genomic studies.

## Author contributions

V.K.S. conceived and coordinated the project. S.M. performed sample collection, DNA-RNA extraction, prepared samples for sequencing, performed long read sequencing, species identification assays, functional annotation of gene sets, and constructed all the figures. M.S.B. and A.C. designed computational framework of the study, and performed all the computational analyses presented in the study. S.M., M.S.B, A.C. and V.K.S. analyzed the data and interpreted the results. S.M. and V.K.S. wrote the first draft of manuscript. S.M., A.C., M.S.B. and V.K.S. wrote and prepared the final manuscript. All the authors have read and approved the final version of the manuscript.

## Acknowledgements

S.M. and A.C. thank Council of Scientific and Industrial Research (CSIR) for fellowship. M.S.B. thanks Ministry of Education, Govt. of India for Prime Minister Research Fellowship (PMRF). The authors also thank the sequencing facilities at Central Instrumentation Facility, IISER Bhopal and the intramural research funds provided by IISER Bhopal.

## Data availability statement

The raw genome and transcriptome reads of *P. emblica* have been submitted in National Center for Biotechnology Information (NCBI).

